# Sleep time, social jetlag and intelligence: biology or work timing?

**DOI:** 10.1101/837443

**Authors:** Péter P. Ujma, Tanja G. Baudson, Róbert Bódizs, Martin Dresler

## Abstract

Sleep-wake patterns show substantial biological determination, but they are also subject to individual choice and societal pressure. Some evidence suggests that high IQ is associated with later sleep patterns. However, t is therefore unclear whether the relationship between IQ and later sleep is due to biological or social effects, such as timing and flexibility of working hours. We investigated the association between habitual sleep timing during work days and work-free days, working time and intelligence in a sample of 1,172 adults. We found no difference in chronotype, and the later sleep timing of high-IQ individuals on work days was fully accounted for by later work start times.

Our results indicate that later sleep timing in those with higher IQs is not due to physiological differences, but rather due to later or more flexible work schedules. Later working times and the resulting lower social jetlag may be one of the reasons why higher IQ is associated with lower prospective morbidity and mortality.

**Statement of significance:** Some evidence shows that higher intelligence is associated with sleep characteristics, but it is unclear if this is because of biological or social mechanisms. We provide evidence for a social mechanism. We found that high IQ individuals indeed sleep later, but only on working days, and this difference is fully accounted for by later work timing. Our evidence is consistent with a view that highly intelligent individuals sleep later because they can afford to, consequently experience lower social jetlag, and this may partially account for better health outcomes.

## Introduction

Sleep-wake patterns depend on genetic effects, social pressure and previously accumulated sleep debt ^1^. Previous research has established a potential link between either sleep-wake timing or chronotype (phase of entrainment to the day-night cycle) and IQ, but it is unclear whether this effect is biological or social – that is, whether more intelligent people are biologically (most likely genetically) predisposed to sleep at a different time, or whether this is a consequence of environmental factors. A large study ^2^ of 15,197 respondents from the Add Health longitudinal study showed a small but significant linear relationship between childhood IQ and self-reported bedtimes and wake-up times between the ages of 18–28 years (*r*=.013–.053). A meta-analysis of the literature up to 2011 ^3^ excluding the Kanazawa et al. study found a similar association between diurnal preference and cognitive ability (morningness: *N*=2,177, *r*=−.042; eveningness *N*=1519, *r*=.075), with no evidence for publication bias. Some recent work, however, found weak positive associations between chronotype and intelligence in school-age children ^4–6^. Additionally, very large genome-wide association studies ^7, 8^ have also revealed that cognitive ability and morningness are in a negative genetic correlation – that is, genetic variants associated with higher morningness are also associated with lower cognitive ability (*r*_g_=−.15−.17). Paradoxically, academic performance, itself moderately correlated with IQ ^9^, is positively correlated with morningness ^3, 10^.

Almost all previous research was conducted with adolescents or young adults, and most studies did not systematically measure work day and free day sleep-wake preferences and lifestyle factors which may influence this relationship. Specifically, it is currently unknown whether the relationship between IQ and sleep-wake timing reflects individual differences in the biological chronotype or rather in social influences (such as different work and school schedules or differences in the use of stimulants or hypnotics). Even genetic studies are not exempt from this, due to the not always causative nature of genetic correlations ^11^.

The goal of our study was to conduct a new study specifying to what extent sleep time differences as a function of IQ are mediated by the different social environments of high-IQ individuals. We assessed the IQ-sleep timing relationship using a sample of working age, high-IQ adults and age- and sex-matched controls with data about both work day and free day sleep timings and work schedules. We compared the self-reported sleep-wake patterns of adult German Mensa (IQ>130) members to an age- and sex-matched random sample of the MCTQ database^12^ (*N*_total_ =1172, mean age: 38.2 years). In doing so, we investigated differences in sleep timing on work days and weekends separately in order to separate the preferred chronotype from the effects of socially pressure on sleep-wake timing ^1^, and specifically investigated the role of work schedules and additional lifestyle variables.

## Methods

### Sample

We recruited 586 members of the German Mensa Society, an organization where a standardized IQ test score at least two standard deviations above the mean (IQ>130) assessed by trained testers and evaluated by a professional psychologist is a prerequisite for membership. As control, we selected a random sample of the same number (*N*=586) age- and sex-matched controls from the main MCTQ database ^12, 13^. There were 232 females and 354 males in each subgroup with a mean age of 38.2 years (see Table 2 for more details). The study was approved by the ethics committee of the University of Munich and all participants gave informed consent accordingly.

### Materials

All participants filled out the MCTQ questionnaire ^13^, which contains questions about basic sociodemographic data (age, sex, place of residence, height and weight), other occupational and behavioral data (number of work days per week, working hours, light exposure on work and free days) as well as about sleep habits (bedtime, time of lights-off, time of falling asleep, wake-up time and the habitual time of getting out of bed) for work days and free days, respectively. From the responses, sleep duration and midsleep (the mid-point between sleep onset and sleep end) on work and work-free days, social jetlag (absolute midsleep difference between work days and free days), weekly sleep deprivation (the difference between the weekly average sleep duration and work day sleep duration, multiplied by the number of work days) and chronotype-proxy MSFsc (free-day midsleep corrected for sleep debt accumulated during the work week) were computed. We also collected information on work start times, commute time, and the consumption of legal substances acting on the central nervous system (i.e., cigarettes, alcohol, coffee, other caffeinated beverages, and hypnotics).

### Analyses

We calculated the differences between Mensa members and matched controls using independent-sample t-tests. Multiple comparisons were controlled for using the Benjamini-Hochberg method of false detection rate (FDR) correction ^14^. IBM SPSS Statistics Version 20 was used to test the confounding effects of extraneous variables on Mensa-control differences in sleep-wake timing with hierarchical linear regression models. The absence of missing data in the MCTQ was not a prerequisite for inclusion in all analyses. For this reason, missing data were found in some subjects, and the final sample sizes for each variable are reported in the Results and Table 1.

**Table 1.**
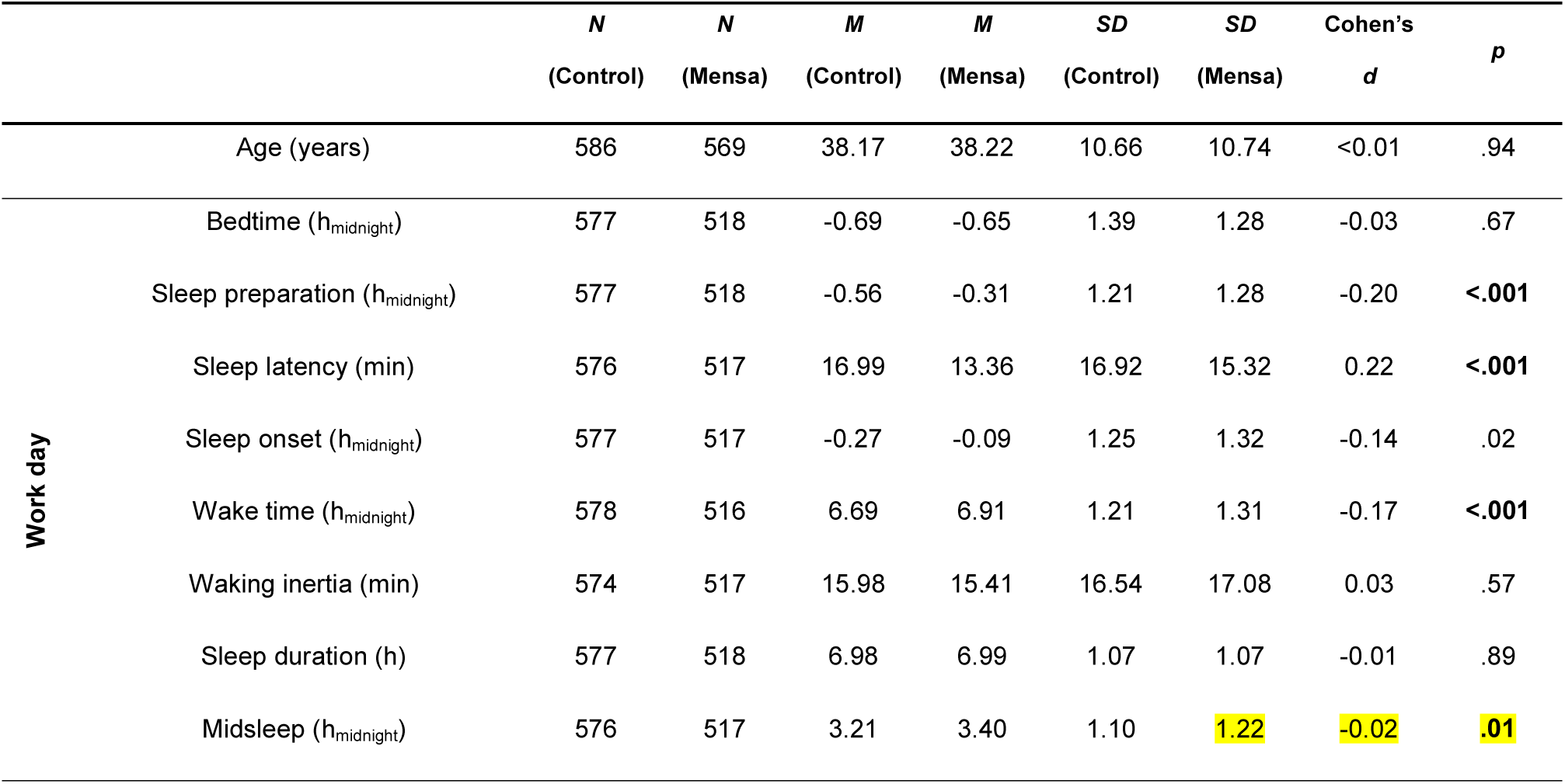
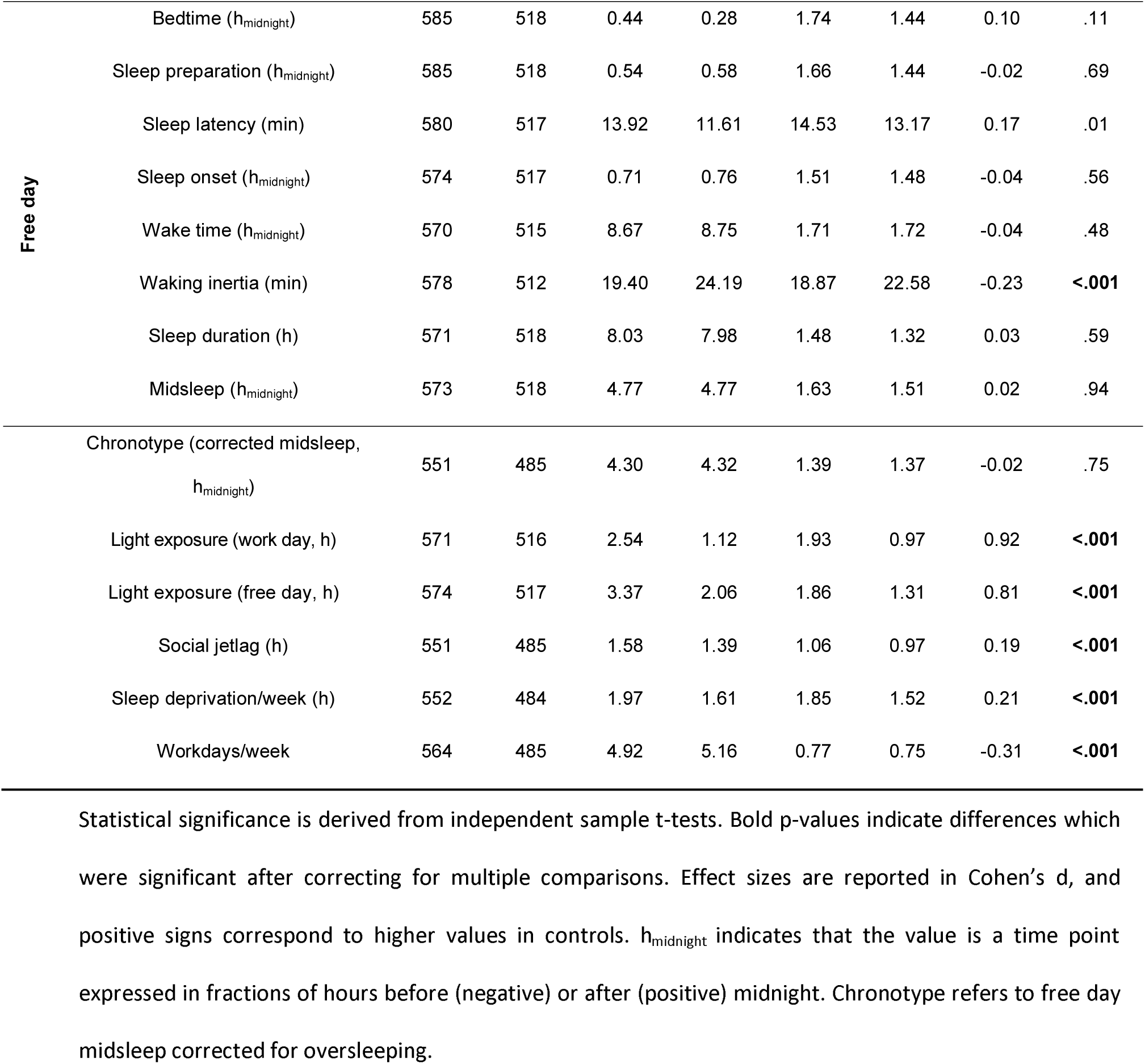
A comparison of Mensa and control samples in terms of chronotype-related and miscellaneous variables.

## Results

We compared the self-reported sleep-wake timing of adult German Mensa (IQ>130) members to an age- and sex-matched random sample of the MCTQ database ^12^ (*N*_total_ =1172). We investigated differences in sleep timing on work day and weekends as well as chronotype (free-day midsleep corrected for oversleeping, MSFsc) separately in order to separate chronotypes from social effects ^1^ and used potential confounding lifestyle variables (Table 3) in hierarchical regression models to test whether they account for subgroup differences.

Mensa members were characterized by later lights-off times, midsleep times and wake-up times as well as shorter sleep latency, but only during work days. During free days, Mensa members still experienced less sleep latency and longer waking inertia (time in bed after waking up), but there was no difference in the main measures in sleep timing. Mensa members suffered from less social jetlag and less weekly sleep deprivation, despite having slightly more work days on average. Importantly, Mensa members had nearly a standard deviation less light exposure, both on free days and working days. Group averages, sample sizes, standard deviations as well as *t*-values and significance levels are reported in Table 1.

So far, our results were in line with some of the literature indicating generally later sleep timing among higher IQ adult participants; however, this difference was limited to work days and not present in the corrected chronotype. Next, we used linear regression models to test whether later work day sleep patterns were accounted for by non-endogenous variables, including workplace characteristics (work start and commute duration) and substance use (consumption of alcohol, cigarettes, caffeine, and hypnotics). Data on all variables were available for 480 Mensa members and 271 controls. Supplementary Table S1 reports descriptive statistics for all covariates by subgroup and Table 2 reports subgroup differences. Supplementary Table S2 provides the correlation matrix of covariates.

**Table 2.**
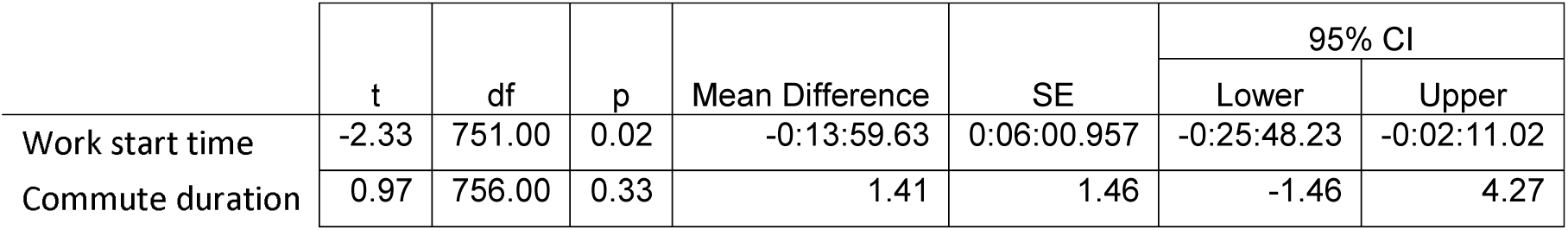
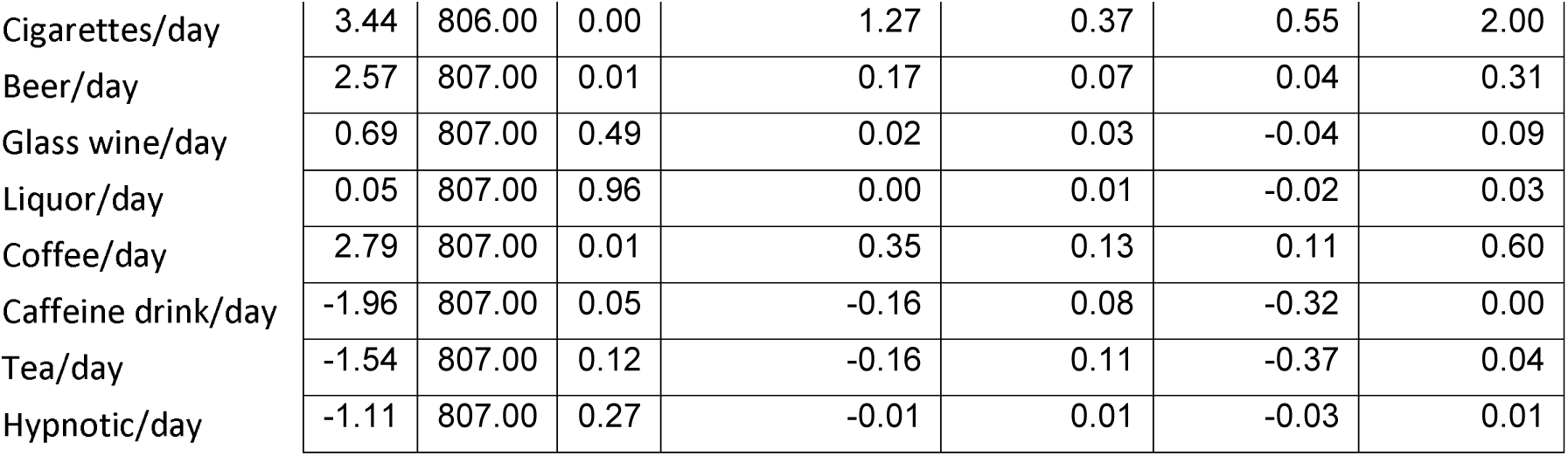
Mensa vs. Control differences in covariates potentially influencing sleep timing. Negative effect sizes indicate higher values in Mensa members. Commute duration is given in minutes. Respondents could provide their consumption of psychoactive substances in monthly, weekly or daily quantities which was always converted to daily values.

|  | t | df | p | Mean Difference | SE | 95% CI |  |
| --- | --- | --- | --- | --- | --- | --- | --- |
|  |  |  |  |  |  | Lower | Upper |
| Work start time | -2.33 | 751.00 | 0.02 | -0:13:59.63 | 0:06:00.957 | -0:25:48.23 | -0:02:11.02 |
| Commute duration | 0.97 | 756.00 | 0.33 | 1.41 | 1.46 | -1.46 | 4.27 |
| Cigarettes/day | 3.44 | 806.00 | 0.00 | 1.27 | 0.37 | 0.55 | 2.00 |
| Beer/day | 2.57 | 807.00 | 0.01 | 0.17 | 0.07 | 0.04 | 0.31 |
| Glass wine/day | 0.69 | 807.00 | 0.49 | 0.02 | 0.03 | -0.04 | 0.09 |
| Liquor/day | 0.05 | 807.00 | 0.96 | 0.00 | 0.01 | -0.02 | 0.03 |
| Coffee/day | 2.79 | 807.00 | 0.01 | 0.35 | 0.13 | 0.11 | 0.60 |
| Caffeine drink/day | -1.96 | 807.00 | 0.05 | -0.16 | 0.08 | -0.32 | 0.00 |
| Tea/day | -1.54 | 807.00 | 0.12 | -0.16 | 0.11 | -0.37 | 0.04 |
| Hypnotic/day | -1.11 | 807.00 | 0.27 | -0.01 | 0.01 | -0.03 | 0.01 |

Work start timing and commute duration were the strongest predictors of work day midsleep timing, and later work start times (*M*_Mensa_=8:35, *SD*=1:18; *M*_Control_=8:21, SD=1:21, values indicating hours and minutes) among Mensa members alone fully mediated the association between Mensa membership and work day midsleep timing (Table 3).

**Table 3.** Results of linear regression models with work day midsleep timing as the dependent variable.

| Model | Independents | $\beta$ | $p$ |
| --- | --- | --- | --- |
| 1 | Mensa | .096 | .009 |
| 2 | Mensa | .036 | .204 |
|  | Work start time | .626 | <.001 |
|  | Commute duration | -.150 | <.001 |
| 3 | Mensa | .044 | .116 |
|  | Work start time | .630 | <.001 |
|  | Commute duration | -.143 | <.001 |
|  | Cigarettes/day | .094 | .002 |
|  | Beer/day | -.010 | .732 |
|  | Glass wine/day | -.023 | .426 |
|  | Liquor/day | .032 | .252 |
|  | Coffee/day | .014 | .624 |
|  | Caffeine drink/day | .071 | .014 |
|  | Tea/day | -.023 | .404 |
|  | Hypnotics/day | .023 | .403 |

Positive standardized regression coefficients for “Mensa” indicate later midsleep timing in Mensa members. Model 3 incorporates the self-reported number of various alcoholic beverages, coffee and other caffeinated beverages, cigarettes and hypnotics medication consumed on a typical day.

In sum, we found no statistically significant differences between the preferred sleep timings of highly intelligent individuals and matched controls. Sleep timing differences on work days were fully accounted for by the later work schedules of Mensa members.

## Discussion

We assessed sleep timing differences between highly intelligent subjects and matched controls. We did find later midsleep timing and waking time in Mensa members, but this was limited to work days, and even then, it was fully accounted for by the later work start times. We found no subgroup difference in chronotype. Thus, it appears that earlier sleep timing in those with lower IQs is not due to physiological differences, but rather a function of earlier work schedules, and consequently only appears in individuals who are old enough to work. Much higher levels of both working day and free day light exposure was found in control participants. While our database does not allow the empirical investigation of the causes of this, we speculate that since IQ is associated with job prestige^15^, extremely intelligent individuals may be overrepresented in jobs requiring clerical office work and underrepresented in outdoor jobs, such as agriculture or construction. Free-day differences might be explained by the well-documented^16–19^ concentration of better educated or more intelligent individuals in urban areas, where outdoors activities are potentially less available or less attractive. In any case, despite large differences in self-reported light-exposure, subgroup differences in sleep timing are modest and limited to work days.

Our findings are partially at odds with some of the previous literature, which generally reported negative associations in young adults ^2, 3^, including pleiotropic genetic effects working in opposite directions ^20^. This discrepancy may be accounted for by two study design features which were uniform in most of the previous literature but differed in ours. First, all previous studies used adolescent or very young adult (≤25 years old) samples, while in ours, participants (38 years on average) were closer to middle age. Second, all studies referenced in the Preckel et al. meta-analysis used questionnaires (typically the MEQ, the LOCI [Lark-Owl Chronotype Indicator], or the CSM) to obtain a single measure of morningness and/or eveningness. These questionnaires – while containing questions about both preferred and usual activity patterns and being correlated with other measures such as the MCTQ ^21^ – do not strictly separate work day and weekend habits, in contrast to the MCTQ used in our study. Combining work day and weekend sleep timing into a general chronotype measure offers a limited opportunity to distinguish individual differences due to biological preferences (most prominent on weekends) and those due to societal pressure (most prominent on work days). Notably, the study of Kanazawa et al ^2^, which enquired about work day and free day bedtimes separately, is an exception to this. This contradiction might arise from several factors, including self-report inaccuracy (as wake times were quite late, on average 10:00 AM or later on weekends and later than 7:00 AM on work days, with some inaccuracies such as frequent 12:00 PM [noon] bedtimes manually corrected by the authors) and age-specific effects as all respondents were very young (<30 years old). Further possibilities are that IQ-sleep timing associations are culture-specific, and while brighter individuals might indeed prefer sleeping later in the United States, they do not do so in Germany, supporting the role of non-endogenous environmental factors such as differences in societal pressure in creating a later chronotype in brighter individuals. A similar caveat applies to the evidence for a negative genetic correlation (*r*_g_=0.15-.17) between morningness and IQ ^20^. The genetic correlation between morningness and IQ is based on summary genetic association data from a chronotype GWAS ^7^, in which morningness was assessed by respondents’ self-report of being a “morning/evening person” on a subjective Likert-type scale. Genetic correlations do not necessarily implicate a common biological mechanism. They may arise from biological pleiotropy if the same genes are implicated in the actual biological processes regulating two phenotypes, but they may also arise from mediated pleiotropy, in which one trait is caused by an environmental influence which in turn is caused by the other, heritable trait ^22^. Our results suggest that the negative genetic correlation between morningness and IQ may be a case of mediated rather than biological pleiotropy: genetic effects resulting in higher IQs do not cause lower morningness because these traits share their biological underpinnings, but because heritable high intelligence exposes individuals to environments – such as more permissive work schedules – in which an early sleep-wake pattern is less preferred. Future studies using Mendelian Randomization (MR) on genetic data may test this hypothesis, which would be supported by a unidirectional causal path from IQ-related genetic variants to sleep timing. We are aware of one such study ^23^, which found minimal causal influence of education on chronotype using an education polygenic score which is also associated with intelligence ^24^.

Our study suffers from a number of limitations. First, due to its cross-sectional design we had a limited ability to assess causation. Because IQ is stable throughout the lifetime ^25^ and highly heritable ^26^, and employees are usually unable to freely choose when their work day starts, we believe that it is more likely that the correlation between IQ and sleep timing as well as work start and sleep timing reflects the effect of the former on the latter than vice versa, but only longitudinal designs could test this reliably. Second, our sample may have been underpowered to detect very small associations between IQ and free day sleep timing or chronotype. Third, we did not have IQ data available from our control sample, therefore, we are unable to perform fine-grained analyses within the normal IQ range or quantitatively assess the effects of range extension. Because participation in the control group required being interested in and filling out an internet-based questionnaire, it is likely that the average IQ of our control group was above the population average. Fourth, it must be emphasized that the relationship between sleep timing and intelligence was – in line with previous results ^3^ – modest, amounting to a weekday midsleep timing about 14 minutes later and approximately 19 minutes less social jetlag per an approximately 1-2 SD IQ difference, assuming a Mensa mean not much higher than 130 and a control group mean above 100 but not exceeding 115.

Overall, our analysis of an adult sample suggests that highly intelligent individuals do not have a biological preference for a later chronotype: while they do go to bed, sleep, and wake up later on work days, this effect is fully mediated by later working times and not present on free days or in the corrected chronotype. Highly intelligent individuals also suffer lower social jetlag. Thus, the effect of intelligence on chronotype seems to have less to do with biological differences than with different work-related environmental pressures on sleep schedules. Eveningness and specifically social jetlag has negative effects on overall morbidity and mortality, including leading causes of death ^27, 28^, high IQ, however, is associated with a lower chance of all-cause mortality ^29^ and morbidity ^30, 31^. Our results suggest that reduced social jetlag in those with late chronotypes due to later or more permissive work schedules might be one of the factors mediating the protective role of high IQ against morbidity and mortality. While the mean difference in social jetlag between Mensa members and controls was modest, amounting to about 19 minutes compared to a pooled average of about 1.5 hours with a standard deviation of 1 hours, a mean difference of this size can generate very large differences at the extreme values of the distributions where pathogenic quantities of social jetlag are likely to be found. Assuming a normal distribution and extrapolating the means and standard deviations reported in Table 1 to the general population it can be calculated that about 9% of individuals with an IQ equal to the control group – likely already above the population mean – can be expected to have a social jetlag over 3 hours and 1.1% over 4 hours, compared to 4.8% and 0.3% for individuals with IQ>130, resulting in a substantially elevated disease risk as a function of lower IQ. Because our results indicate that these risk differences are the result of lifestyle factors rather than biological differences, policies enabling a better alignment of desired and actual work schedules could be beneficial to individuals not possessing high intelligence.

## Acknowledgements

We thank Mensa in Deutschland e.V. (MinD) for their help in recruiting high-IQ participants. We further thank Celine Vetter and Till Roenneberg for their help with data acquisition and their valuable comments on the manuscript.

