## Supplementary Table S1 for "Sleep time, social jetlag and intelligence: biology or work timing?"

**Supplementary table S1. Descriptive statistics for the covariates described in Table 3.**

|  |  | N | Minimum | Maximum | Mean | SD |
| --- | --- | --- | --- | --- | --- | --- |
| Mensa members | Work start time | 481 | 04:00:00 | 15:00:00 | 08:35:23 | 01:18:15 |
|  | Commute duration | 483 | 0.0 | 120.0 | 22.8 | 19.6 |
|  | Cigarettes/day | 515 | 0.0 | 40.0 | 1.2 | 4.6 |
|  | Beer/day | 515 | 0.0 | 12.6 | 0.3 | 0.8 |
|  | Glass wine/day | 515 | 0.0 | 4.3 | 0.2 | 0.5 |
|  | Liquor/day | 515 | 0.0 | 2.0 | 0.1 | 0.2 |
|  | Coffee/day | 515 | 0.0 | 13.6 | 1.5 | 1.8 |
|  | Caffeine drink/day | 515 | 0.0 | 14.3 | 0.4 | 1.3 |
|  | Tea/day | 515 | 0.0 | 14.3 | 0.6 | 1.6 |
|  | Hypnotic/day | 515 | 0.0 | 1.0 | 0.0 | 0.2 |
| Controls | Work start time | 272 | 5:30:00 | 16:00:00 | 8:21:23 | 1:21:06 |
|  | Commute duration | 275 | 0.0 | 100.0 | 24.2 | 18.8 |
|  | Cigarettes/day | 293 | 0.0 | 28.6 | 2.5 | 5.8 |
|  | Beer/day | 294 | 0.0 | 14.3 | 0.4 | 1.1 |
|  | Glass wine/day | 294 | 0.0 | 2.9 | 0.3 | 0.4 |
|  | Liquor/day | 294 | 0.0 | 1.4 | 0.1 | 0.2 |
|  | Coffee/day | 294 | 0.0 | 10.0 | 1.9 | 1.7 |
|  | Caffeine drink/day | 294 | 0.0 | 7.1 | 0.3 | 0.8 |
|  | Tea/day | 294 | 0.0 | 7.1 | 0.5 | 1.0 |
|  | Hypnotic/day | 294 | 0.0 | 1.0 | 0.0 | 0.1 |
