## Supplementary Table S2 for "Sleep time, social jetlag and intelligence: biology or work timing?"

**Supplementary table S2. Pearson’s correlation coefficients between the covariates described in Table 3.**

|  | | Work start time | Commute duration | Cigarettes/day | Beer/day | Glass wine/day | Liquor/day | Coffee/day | Caffeine drink/day | Tea/day | Hypnotic/day |
| --- | --- | --- | --- | --- | --- | --- | --- | --- | --- | --- | --- |
| Work start time | r | 1 | -.051 | -.063 | -.007 | .021 | -.010 | -0.105 | .046 | .065 | .069 |
|  | p |  | .162 | .087 | .854 | .560 | .793 | .004 | .209 | .077 | .060 |
|  | N | 753 | 753 | 746 | 747 | 747 | 747 | 747 | 747 | 747 | 747 |
| Commute duration | r | -.051 | 1 | -.006 | -.023 | -.035 | -.041 | -.002 | -0.072 | -.010 | -.005 |
|  | p | .162 |  | .863 | .528 | .338 | .262 | .953 | .048 | .779 | .892 |
|  | N | 753 | 758 | 751 | 752 | 752 | 752 | 752 | 752 | 752 | 752 |
| Cigarettes/day | r | -.063 | -.006 | 1 | 0.331 | .062 | .010 | 0.234 | 0.136 | -.065 | -.011 |
|  | p | .087 | .863 |  | .000 | .079 | .768 | .000 | .000 | .066 | .754 |
|  | N | 746 | 751 | 808 | 808 | 808 | 808 | 808 | 808 | 808 | 808 |
| Beer/day | r | -.007 | -.023 | 0.331 | 1 | 0.079 | 0.174 | 0.224 | -.039 | -.021 | .019 |
|  | p | .854 | .528 | .000 |  | .025 | .000 | .000 | .262 | .559 | .587 |
|  | N | 747 | 752 | 808 | 809 | 809 | 809 | 809 | 809 | 809 | 809 |
| Glass wine/day | r | .021 | -.035 | .062 | .079^*^ | 1 | .062 | 0.186 | -0.104 | 0.091 | .039 |
|  | p | .560 | .338 | .079 | .025 |  | .079 | .000 | .003 | .010 | .270 |
|  | N | 747 | 752 | 808 | 809 | 809 | 809 | 809 | 809 | 809 | 809 |
| Liquor/day | r | -.010 | -.041 | .010 | 0.174 | .062 | 1 | .019 | -.033 | -.024 | -.005 |
|  | p | .793 | .262 | .768 | .000 | .079 |  | .592 | .354 | .500 | .894 |
|  | N | 747 | 752 | 808 | 809 | 809 | 809 | 809 | 809 | 809 | 809 |
| Coffee/day | r | -0.105 | -.002 | 0.234 | 0.224 | 0.186 | .019 | 1 | -0.117 | -.055 | -0.082 |
|  | p | .004 | .953 | .000 | .000 | .000 | .592 |  | .001 | .117 | .020 |
|  | N | 747 | 752 | 808 | 809 | 809 | 809 | 809 | 809 | 809 | 809 |
| Caffeine drink/day | r | .046 | -0.072 | .136 | -.039 | -0.104 | -.033 | -0.117 | 1 | -.044 | .043 |
|  | p | .209 | .048 | .000 | .262 | .003 | .354 | .001 |  | .215 | .225 |
|  | N | 747 | 752 | 808 | 809 | 809 | 809 | 809 | 809 | 809 | 809 |
| Tea/day | r | .065 | -.010 | -.065 | -.021 | .091^**^ | -.024 | -.055 | -.044 | 1 | -.012 |
|  | p | .077 | .779 | .066 | .559 | .010 | .500 | .117 | .215 |  | .729 |
|  | N | 747 | 752 | 808 | 809 | 809 | 809 | 809 | 809 | 809 | 809 |
| Hypnotic/day | r | .069 | -.005 | -.011 | .019 | .039 | -.005 | -0.082 | .043 | -.012 | 1 |
|  | p | .060 | .892 | .754 | .587 | .270 | .894 | .020 | .225 | .729 |  |
|  | N | 747 | 752 | 808 | 809 | 809 | 809 | 809 | 809 | 809 | 809 |
